# Evaluation of vectors for gene expression in *Pseudovibrio* marine bacteria

**DOI:** 10.1101/2024.08.01.606211

**Authors:** Yitao Dai, Alessandra S. Eustáquio

**Affiliations:** Department of Pharmaceutical Sciences and Center for Biomolecular Sciences, College of Pharmacy, University of Illinois at Chicago, Chicago, IL, USA

**Keywords:** plasmid vectors, *Pseudovibrio*, conjugation, electroporation, fluorescent protein, swarming

## Abstract

α-Proteobacteria belonging to the *Pseudovibrio* genus have been isolated from different marine organisms including marine sponges, corals and algae. This genus was first described in 2004 and has since garnered attention due to the potential ecological relevance and biotechnological application of its metabolites. For instance, we recently reported specialized metabolites we named pseudovibriamides from *Pseudovibrio brasiliensis* Ab134. The pseudovibriamide encoding *ppp* gene cluster is found in two thirds of *Pseudovibrio* genomes. Pseudovibriamides coordinate motility and biofilm formation, behaviors that are known to be important for host colonization. Although we previously established reverse genetics methods to delete genes via homologous recombination, no self-replicative vectors have been reported for *Pseudovibrio*. We show that plasmid vectors containing two different broad-host-range replicons, RSF1010 and pBBR1, can be used in *P. brasiliensis*. The efficiency of vector transfer by electroporation averaged ∼ 3 × 10^3^ CFU/µg plasmid DNA whereas the conjugation frequency from *E. coli* ranged from 10^-3^ to 10^-6^. We then tested the vectors for fluorescent protein expression and consequent labeling, which allowed us to observe their effects on swarming motility and to compare plasmid stability. This study expands the genetic toolbox available for *Pseudovibrio* which is expected to enable future ecological and biotechnological studies.

**Importance:** The genus *Pseudovibrio* of α-Proteobacteria has consistently been isolated from marine sponges and other marine organisms such as corals and algae. *Pseudovibrio* bacteria are a source of antibiotics and other secondary metabolites with the potential to be developed into pharmaceuticals. Moreover, the secondary metabolites they produce are important for their physiology and for interactions with other organisms. Here we expand the genetic tool box available for *Pseudovibrio* bacteria by establishing self-replicative vectors that can be used for the expression of e.g., fluorescent proteins. The availability of genetic tools is important to enable us to explore the emerging ecological and biotechnological potential of *Pseudovibrio* bacteria.

## Introduction

The genus *Pseudovibrio* of α-Proteobacteria has consistently been isolated from marine sponges, in addition to corals and algae, among other marine organisms [1]. *Pseudovibrio* has been hypothesized to contribute to marine sponge health because it has been isolated only from healthy sponges, it produces antibiotics that prevent the growth of sponge pathogens, and evidence for vertical transmission has been provided [1]. Beyond sponges, *Pseudovibrio* has also been shown to induce coral larvae settlement behavior [2].

Genetics tools are important to facilitate studies on *Pseudovibrio* biology and for future biotechnological applications. We recently discovered specialized metabolites we named pseudovibriamides (**Figure 1**) from *Pseudovibrio brasiliensis* Ab134 [3]. The pseudovibriamide encoding gene cluster (termed *ppp*) is present in approximately two thirds of *Pseudovibrio* genomes [3]. Reverse genetics methods we established (gene knockout via homologous recombination) helped us discover a link between pseudovibriamides and *Pseudovibrio* collective behaviors, that is, flagella-mediated motility and biofilm formation [3]. Motility and biofilm formation are known to be important for bacterial survival and for host colonization [4].

**Figure 1.**
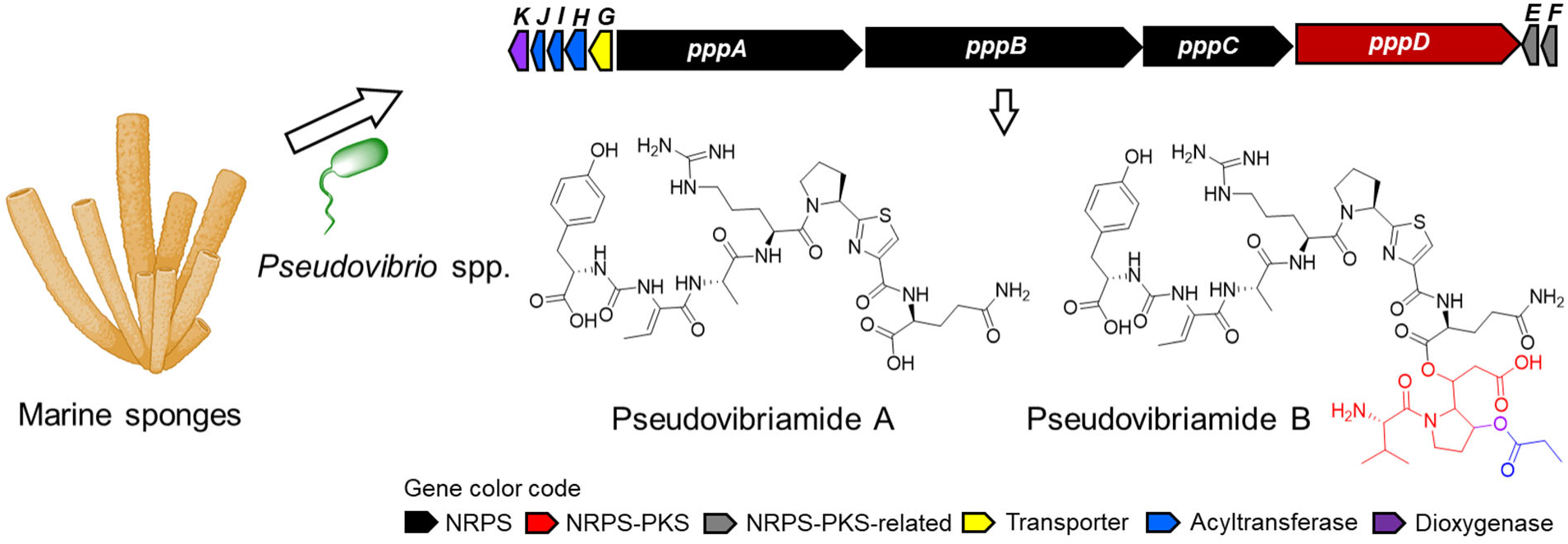
Pseudovibriamides from *Pseudovibrio* bacteria isolated from marine sponges. Structures and encoding gene cluster. Pseudovibriamides were isolated from *Pseudovibrio brasiliensis* Ab134 [3]. The *ppp* gene cluster is present in 2/3 of *Pseudovibrio* genomes isolated from various marine sponges including *Arenosclera brasiliensis*, *Axinella dissimilis*, *Spongia officinalis*, *Polymastia penicillus*, [3] and *Aplysina aerophoba* [5]. NRPS, nonribosomal peptide synthetase, PKS, polyketide synthase. The sponge icon is from BioRender.com.

Although methods for gene inactivation are available [3], no self-replicative vectors that could aid with gene expression have been reported for *Pseudovibrio*. Here we compared broad-host range self-replicative vectors and identified those that can be used with *P. brasiliensis* to express fluorescent proteins. We also report conjugation and electroporation protocols and compare vector stability and their effect on swarming motility.

## Results

### Establishing broad-host range vectors for *P. brasiliensis* Ab134

We started by obtaining and testing plasmid vectors containing two different broad-host-range replicons, RSF1010 and pBBR1. The corresponding plasmids pAM4891 [6], pSEVA234M, and pSEVA237R_Pem7 [7] were transferred into *P. brasiliensis* Ab134 by conjugation from *E. coli* and obtained clones were confirmed by plasmid extraction, restriction digestion and gel electrophoresis (**SI Figure S1**). We then used short-read Illumina sequencing of total DNA to estimate relative plasmid copy number by mapping the obtained Illumina reads to the Ab134 reference genome [3]. We have previously shown that Illumina sequencing is comparable to quantitative PCR for relative plasmid copy number determination [8]. Others have also used sequencing instead of quantitative PCR for this purpose [9]. The vector coverage normalized to the chromosome coverage indicated that RSF1010 is of medium copy number (49) and that pBBR1 is of high copy number (148 to 210) in *P. brasiliensis* Ab134 (**Table 1**).

**Table 1.**
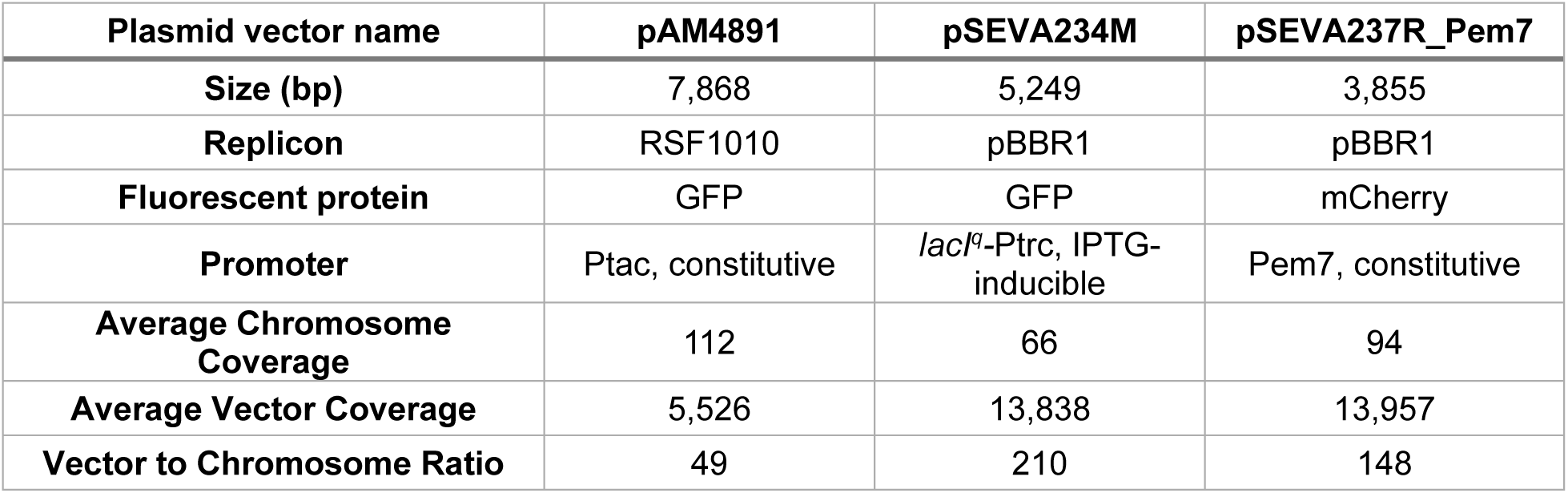
Normalized plasmid vector copy number in *P. brasiliensis* Ab134. Data obtained from Illumina sequencing of total DNA.

Growth comparison in liquid cultures showed that *P. brasiliensis* Ab134 carrying pAM4891 with the RSF1010 replicon had comparable growth in terms of total area under the growth curve to the wild type but a ∼40% longer lag phase (**Figure 2**). In contrast, Ab134 carrying pSEVA234M and pSEVA237R_Pem7, both with the same pBBR1 replicon, showed distinct growth phenotypes, with pSEVA234M displaying a growth defect. Although the lag phase was not affected (**Figure 2C**), the final OD_600_ and the area under the growth curve were significantly reduced (**Figure 2AB**; **SI Table S2**). The marked growth defect in Ab134 carrying pSEVA234M was consistent across independent clones (**SI Figure S2**). Although it has been shown that high plasmid copy number can result in growth defects [10, 11], Ab134 carrying pSEVA237R_Pem7 with the same pBBR1 replicon did not exhibit this defect (**Figure 2** and **SI Figure S2**). This discrepancy could be due to the higher plasmid copy number of pSEVA234M (210) compared to pSEVA237R_Pem7 (148) (**Table 1**). Another possibility is that other components within the pSEVA234M vector, such as repressor-promoter *lacI^q^-*Ptrc, may be responsible. The mutation in the promoter of the *lacI* repressor gene (*lacI^q^*) results in higher levels of LacI within cells [12]. The overexpression of foreign genes can lead to decreased growth rate as demonstrated with *E. coli* [13]. Thus, the combination of high plasmid copy number and high expression of the *lacI* repressor gene from pSEVA234M could be the underlying cause of the observed growth defect.

**Figure 2.**
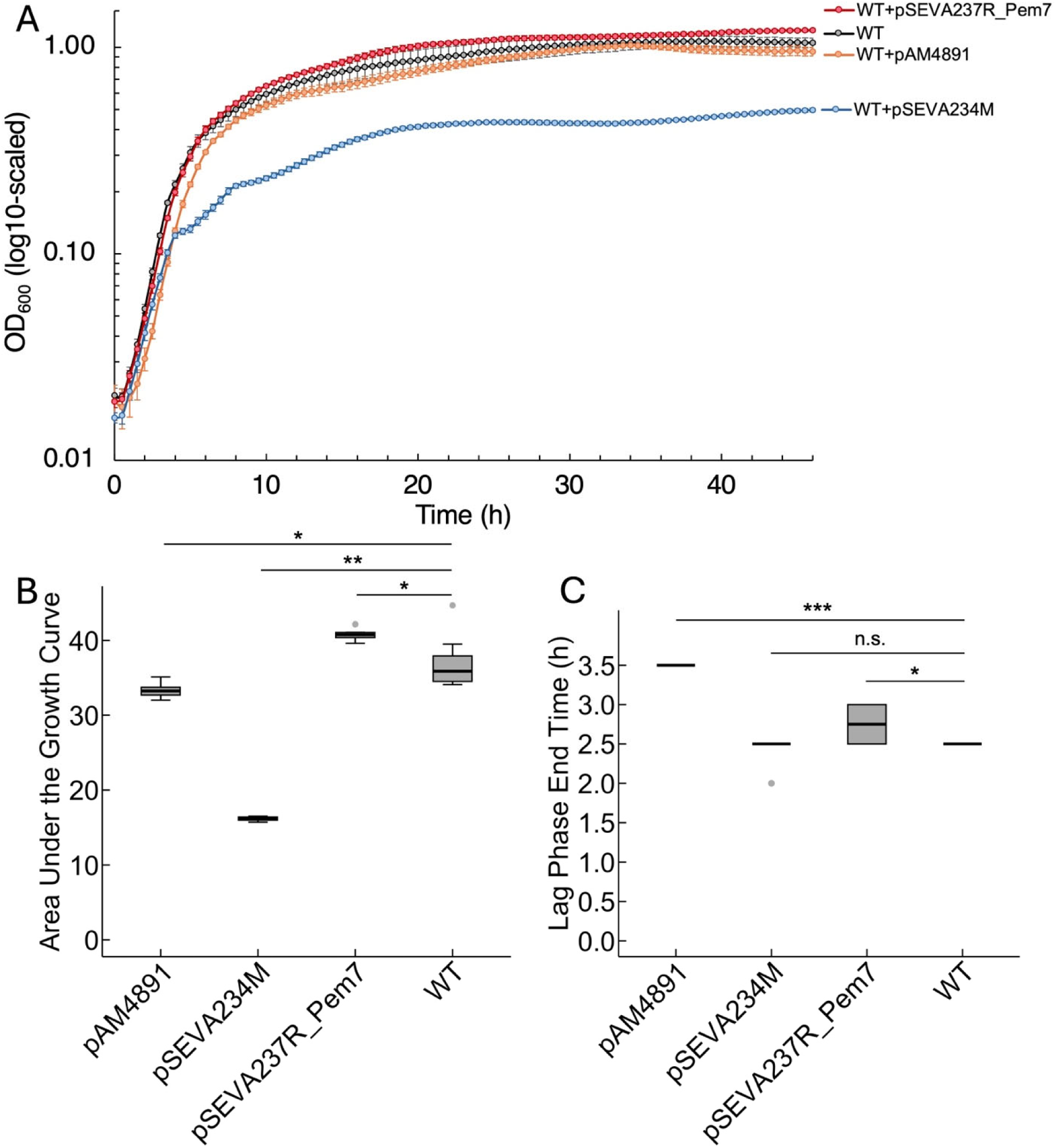
Growth of *P. brasiliensis* Ab134 containing different vectors. (**A**) Growth in liquid cultures was measured by optical density at 600 nm (OD_600_). *N*=10. Error bars indicate standard deviation. The y-axis is scaled using a base-10 logarithmic (log_10_) transformation. WT, *P. brasiliensis* Ab134 wild type. See Table 1 for vector details. Comparison of (**B**) the area under the growth curve (not log-scaled) and (**C**) the lag phase of *P. brasiliensis* Ab134 containing different vectors using box and whisker plots. *N*=10. Error bars indicate the 75^th^ and 25^th^ percentiles ± 1.5× interquartile range. Grey dots indicate outliers. Two-Sample *t* test was implemented for statistical analyses. *, *P* ≤ 0.01; **, *P* ≤ 10^-9^; ***, *P* ≤ 1/ ∞; n.s., not significant.

Thus, we moved on with pAM4891 and pSEVA237R_Pem7 because these two plasmids had the least effect on growth.

### Plasmid pSEVA237R_Pem7 outperforms pAM4891 in conjugation but not electroporation

We next evaluated the DNA transfer efficiency of pAM4891 and pSEVA237R_Pem7 into *P. brasiliensis* Ab134 by electroporation and by conjugation from *E. coli* S17-1 which are common methods to introduce plasmid DNA into bacteria. Both pAM4891 and pSEVA237R_Pem7 constitutively express a fluorescent protein gene (*gfp* and *mCherry*, respectively), allowing quick fluorescence-based screening of positive clones.

The electroporation efficiency was comparable between plasmid pAM4891 (3,295 ± 271 CFU per µg plasmid DNA) and pSEVA237R_Pem7 (2,776 ± 495 CFU per µg plasmid DNA), with no statistically significant difference (**Figure 3A** and **SI Table S3**). In contrast, the conjugation frequency of pSEVA237R_Pem7 (2.22 × 10^-3^ ± 6.69 × 10^-4^) was significantly higher than that of pAM4891 (3.86 × 10^-6^ ± 3.34 × 10^-7^) (**Figure 3B** and **SI Table S4**).

**Figure 3.**
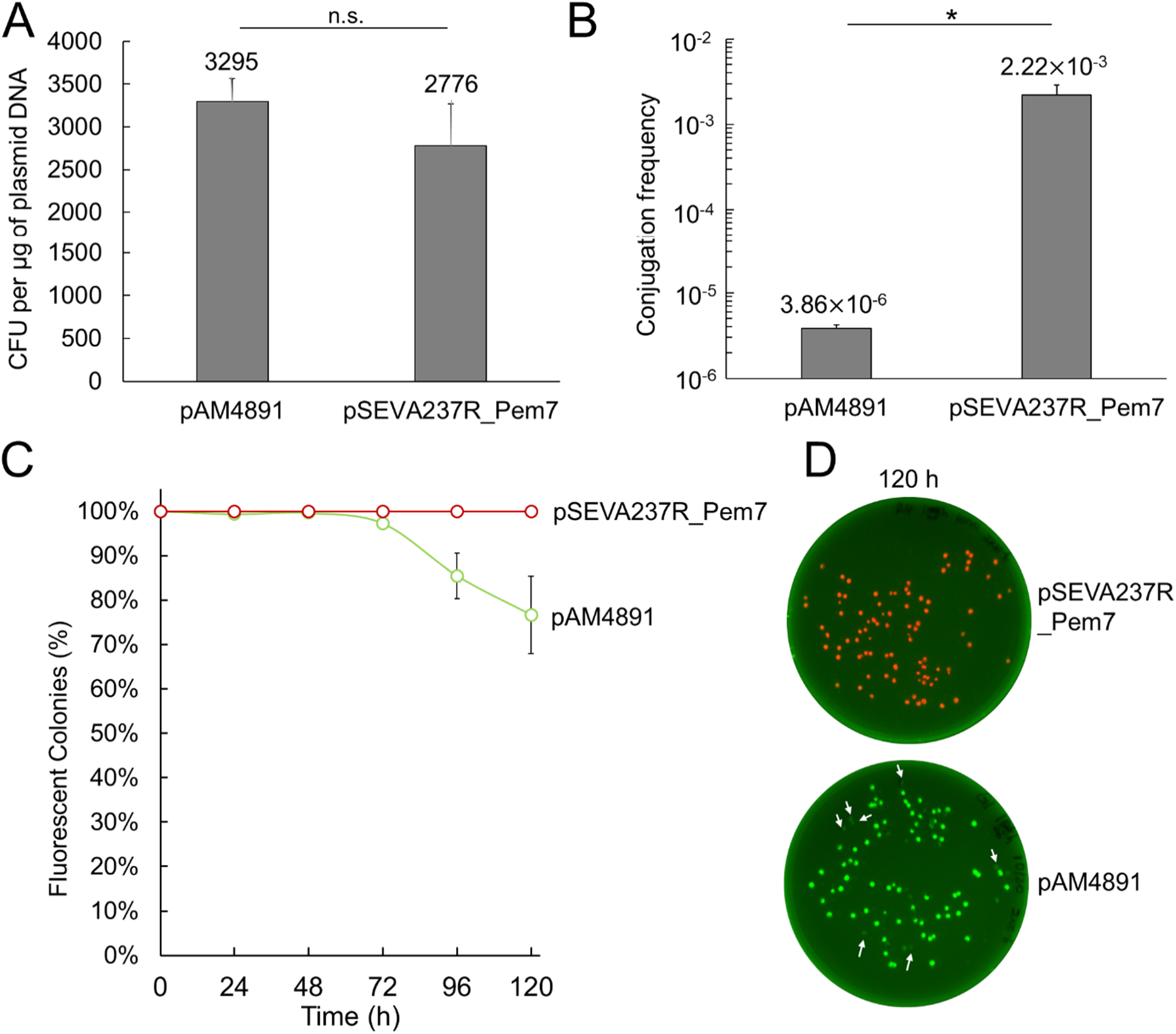
DNA transfer efficiency and stability of *P. brasiliensis* Ab134 containing pAM4891 and pSEVA237R_Pem7. (**A**) Comparison of the electroporation efficiency of pAM4891 and pSEVA237R_Pem7 into *P. brasiliensis* Ab134. The electroporation efficiency represents the number of colonies formed per microgram of plasmid DNA used. The assay was performed in triplicates (**SI Table S3**). Two-sample *t* test was implemented for statistical analyses. n.s., not significant. (**B**) Comparison of conjugation efficiency for plasmid transfer (pAM4891 and pSEVA237R_Pem7) into *P. brasiliensis* Ab134 from *E. coli* S17-1. The conjugation efficiency is defined as the ratio between the colony-forming units (CFU) of transconjugants per milliliter and the CFU of *P. brasiliensis* Ab134 recipients per milliliter. The assay was performed in triplicates (**SI Table S4**). Two-sample *t* test was implemented for statistical analyses. *, *P* ≤ 0.01. The y-axis is scaled using a base-10 logarithmic (log_10_) transformation. (**C**) Comparison of plasmid stability between pAM4891 and pSEVA237R_Pem7 in *P. brasiliensis* Ab134 over 5 passages for a total of 120 hours, with approximately ten generations in each passage (24-hour period). Fluorescence was used as an indicator to quickly assess plasmid retention in the absence of antibiotics. The assay was performed in quadruplicates. (**D**) Representative images of plates at the 120-hour time point showing *P. brasiliensis* Ab134 colonies containing pSEVA237R_Pem7 (red fluorescence), and pAM4891 (green fluorescence), respectively. White arrows indicate colonies that lost fluorescence, indicating plasmid loss.

### Both pAM4891 and pSEVA237R_Pem7 are stable in passage studies up to three days and pSEVA237R_Pem7 up to five days

Another parameter of interest is plasmid stability. Evaluating plasmid stability over generations in the absence of antibiotic selection is advantageous for minimizing the potential unintended effects of antibiotics on phenotypes, such as swarming motility, and metabolism [14, 15] We assessed the stability of pAM4891 and pSEVA237R_Pem7 over five serial passages (24 hours each), totaling 120 hours (**Figure 3C**). Due to the constitutive expression of *gfp* in pAM4891 and *mCherry* in pSEVA237R_Pem7, colonies lacking fluorescence were classified as having lost the (**Figure 3D**). Over 97% of Ab134 cells retained pAM4891 during the first three passages (72 hours), but this retention rate dropped to 85.4% by the fourth passage and to 76.6% by the fifth passage (**Figure 3C** and **SI Table S5**). In contrast, 100% of Ab134 cells retained pSEVA237R_Pem7 across all five passages (120 hours), suggesting significantly higher, long-term plasmid stability for pSEVA237R_Pem7 (**Figure 3C** and **SI Table S5**).

### Plasmid pSEVA237R_Pem7 results in a delayed onset of swarming motility

Self-replicative vectors are useful for gene expression including genetic complementation studies or for the labeling of strains with fluorescent proteins. For instance, when studying the effect of gene deletions on bacterial motility, the plasmid used for genetic complementation should ideally not itself affect motility. Moreover, one of our future goals is to test whether pseudovibriamides are important for host colonization. The strains would need to be labeled with fluorescent proteins, and again, the vector itself should not affect motility.

We have previously shown that a pseudovibriamide-defective mutant strain (Δ*pppA*) displays reduced swarming motility compared to the wild type [3]. Here we investigated whether carrying a self-replicative vector would affect the swarming motility of wild type and Δ*pppA* strains, and whether fluorescent proteins could be detected without selection during the swarming assay.

The swarming motility of strains carrying the pAM4891 vector was comparable to the parent strains, whereas strains carrying the pSEVA237R_Pem7 vector exhibited a delayed onset of the swarming phenotype (**Figure 4A** and **SI Figure S6**). Nevertheless, as for the parent strain, the swarming motility of Δ*pppA* strains carrying either vector was still reduced compared to the wild type carrying the same vectors (**Figure 4A** and **SI Figure S6**). Finally, we were able to detect fluorescent protein expression in these strains after three days even without antibiotic selection, indicating that both vectors are stably maintained over the three-day course of the experiment (**Figure 4B**) consistent with the stability studies (**Figure 3C**).

**Figure 4.**
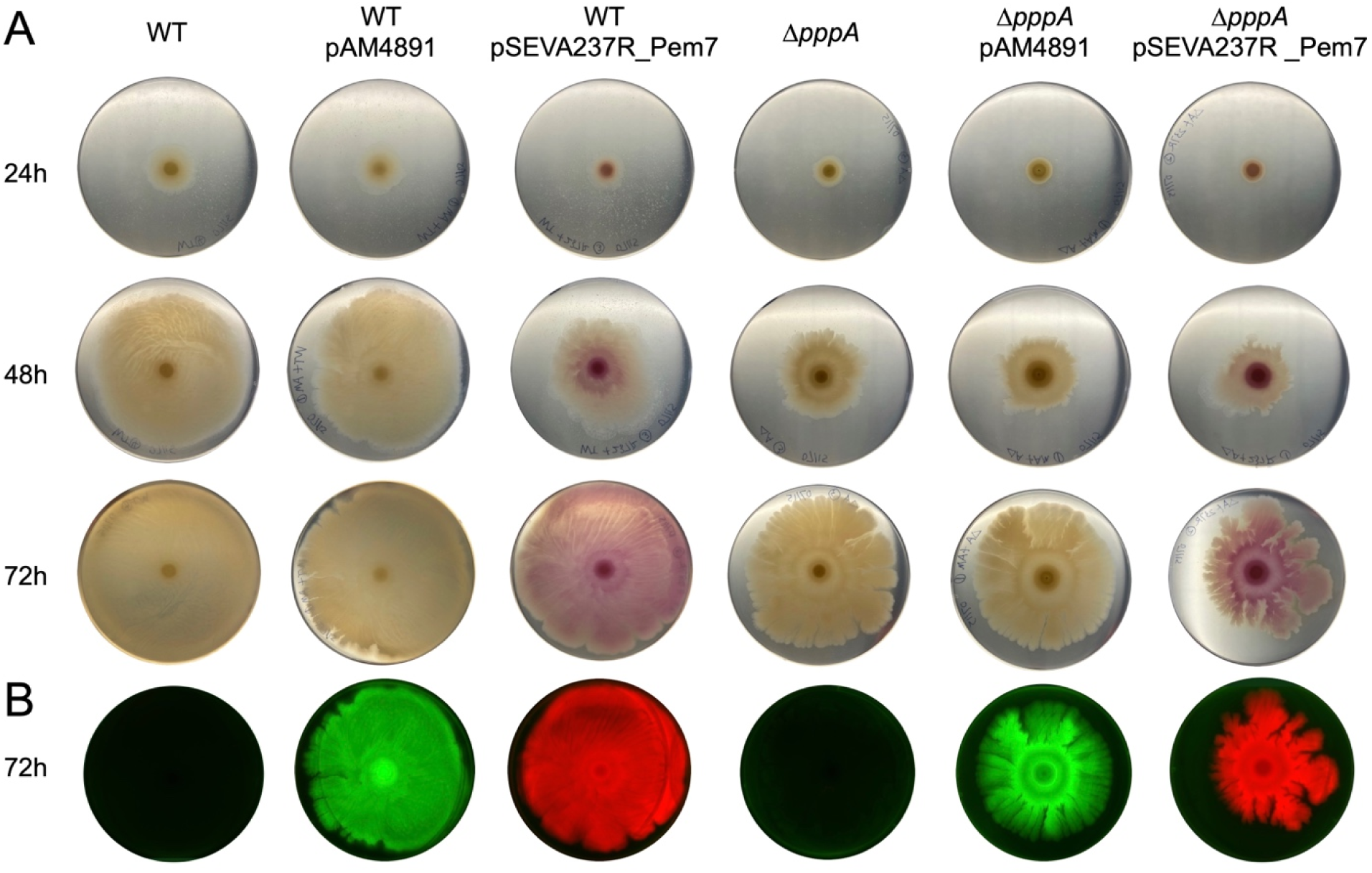
Swarming motility of *P. brasiliensis* Ab134 wild type or Δ*pppA* mutant parent strains compared to strains carrying self-replicative vectors. (**A**) Swarming assays performed on marine broth with 0.5 % Eiken agar. Pictures shown were taken at 24, 48, and 72 hours after inoculation, respectively. The assay was performed in triplicates (**SI Figure S6**) and representative images are shown. (**B**) Fluorescence imaging of swarming plates at 72 hours to show the expression of GFP and mCherry in strains carrying pAM4891 and pSEVA237R_Pem7, respectively.

## Discussion

The genus *Pseudovibrio* was first described in 2004 [16]. Since then, *Pseudovibrio* bacteria have been isolated or detected predominantly from marine sponges and corals worldwide but also from tunicates, flatworms, algae and other sources [1]. In addition to potentially beneficial roles such as supporting biogeochemical cycling and providing nutrients, *Pseudovibrio* has also been proposed to contribute to marine sponge health, and to promote coral larvae settlement [1, 2]. The availability of genetic tools is important to enable us to explore the emerging ecological and biotechnological potential of *Pseudovibrio* bacteria.

Although we have previously established reverse genetics methods for *P. brasiliensis* using a suicide vector [3, 17], no self-replicative vectors for gene expression have been reported for *Pseudovibrio*. Here we showed that vectors containing two distinct, broad-host range replicons can be used in *P. brasiliensis* to express fluorescent proteins GFP and mCherry.

The RSF1010 replicon belongs to the IncQ group of plasmids. The largest of the replicons tested here (5.4 kb) is composed of *oriV*, three replication genes (*repA*, *repB*, and *repC*) and three mobilization proteins (*mobA*, *mobB* and *mobC*). Its broad host range expands to Pseudomonadota (including Alphaproteobacteria), Bacillota, Actinomycetota, and Cyanobacteriota [18, 19]. The reported low copy number of 10 to 12 for wild type RSF1010 in *E. coli* [20] may be explained due to self-mobilization which results in DNA nicking and low yields of extracted DNA. The pAM4891 vector tested here contains a mutation in the RSF1010 *mobA* gene (Y25F) that results in plasmid preparations containing mostly supercoiled DNA in contrast to the mostly nicked wild-type DNA [6], which may explain the higher copy number of 49 we observed (**Table 1**).

The pBBR1 replicon does not belong to IncP, IncQ or IncW groups and may represent a new incompatibility group [21]. It has a broad-host range in Gram-negative bacteria that expands to Pseudomonadota (including Alphaproteobacteria) [21] and Thermodesulfobacteriota [22], and it has also been used in *Microbacterium* (Actinomycetota) [23]. The replicon is composed of *oriV*, one *rep* and one *mob* gene, and as such, it is relatively small [21]. The pBBR1 replicon (1.52 kb) in the SEVA plasmids is smaller than the RSF1010 replicon [7]. It contains *oriV* and *rep*. Instead of the *mob* gene present in the original replicon [21], all SEVA plasmids use *oriT* from RK2 [7]. A medium copy number of 30-40 was reported for *E. coli* and *Bordetella pertussis* [21], whereas a growth defect was reported with some bacteria, including *Pseudomonas putida* [10], *Geobacter sulfurreducens* [22] and *Burkholderia* sp. [8] corresponding to the higher copy number of pBBR1 in these bacteria. Both vectors we tested that contain the pBBR1 replicon resulted in high copy number of 148-210 in *P. brasiliensis* (**Table 1**) but only pSEVA234M led to a growth defect (**Figure 2**). The observed growth defect with pSEVA234M may be due to a combination of higher plasmid copy number of 210 and high expression of the *lacI^q^* repressor gene from pSEVA234M [12, 13]. We observed that the plasmid transfer efficiency of pSEVA237R_Pem7 (3,855 bp) was significantly higher than that of pAM4891 (7,868 bp) in *P. brasiliensis* Ab134 via conjugation, in agreement with previous studies showing that the smaller the plasmid size, the higher the conjugation rate [24]. In contrast there was no significant difference observed with electroporation in terms of CFU/µg of plasmid DNA (**Figure 3AB**). Although electroporation efficiency has also been reported to be inversely correlated with plasmid size, other electroporation parameters contribute to the observed efficiency. For instance, under the low field strength used here (9 kV/cm) the effect of plasmid size in the range we tested has been shown to be negligible [25, 26].

Additionally, conjugation was more efficient than electroporation for introducing both plasmids into *P. brasiliensis* Ab134. Conjugation may bypass DNA restriction-modification systems of recipient strains due to the transfer of single stranded DNA, in contrast to electroporation or other transformation methods where double stranded DNA is uptaken [27–29]. Indeed, we identified a type II restriction-modification system (locus tag: KGB56_19030) in *P. brasiliensis* Ab134 using DefenseFinder [30]. Type II restriction-modification systems utilize a restriction endonuclease that recognizes specific palindromic sequences on DNA and performs double-strand cleavage [31]. Although many type II restriction endonucleases have been shown to cleave single stranded DNA, the formation of a double stranded structure with two-fold symmetry is still required for cleavage, and type II restriction endonucleases exhibit lower specificity for single stranded DNA compared to double-stranded DNA [27, 32, 33] The restriction-modification system of *P. brasiliensis* may be harnessed in the future to improve DNA transfer efficiency further if needed such as for library generation larger than 10^3^ CFU.

Although *P. brasiliensis* carrying pSEVA237R_Pem7 and pAM4891 showed comparable growth to the wild type, their swarming motilities were affected to different extents. Specifically, *P. brasiliensis* carrying pSEVA237R_Pem7, with a higher plasmid copy number compared to pAM4891, exhibited a noticeable reduction in swarming motility (**Figure 4A**). Previous studies have shown that the production of unnecessary proteins, including fluorescent proteins, can impose a metabolic burden on the host, slowing native protein synthesis [34–37]. Thus, the production of mCherry from the high copy number pSEVA237R_Pem7 in *P. brasiliensis* might induce a burden on cellular processes, reducing the availability of resources to produce essential proteins for swarming motility. Finally, plasmid pSEVA237R_Pem7 was stably maintained up to five days of passaging in non-selective media, whereas pAM4891 resulted in observable plasmid loss after three days (**Figure 3CD**).

Based on the combined observations, we suggest pAM4891 for genetic complementation experiments since it resulted in negligible effects on swarming motility and it is stable for three days, a time that is sufficient for swarming assays and pseudovibriamide production. In fact, we used pAM4891 for genetic complementation studies reported during review of this manuscript [38].

In conclusion, given the emerging roles of *Pseudovibrio* as part of sponge, coral and other marine holobionts, and their production of secondary metabolites with biomedical and biotechnological potential [1], the availability of genetic tools is important for biotechnological applications but also to facilitate probing mechanistic questions regarding *Pseudovibrio* biology, microorganism-microorganism and microorganism-host interactions.

## Materials and Methods

### General experimental procedures

Chemicals and enzymes were acquired from Sigma-Aldrich, VWR, Becton Dickinson (BD), and Thermo Fisher Scientific, unless otherwise noted. Restriction enzymes were purchased from New England Biolabs (NEB). Genomic DNA was isolated using GenElute^TM^ Bacterial Genomic DNA Kit (Sigma-Aldrich). ZymoPURE Plasmid Miniprep kits were used to extract plasmid DNA. Whole plasmid sequencing by Primordium labs was performed as quality control to confirm that plasmid vector sequences were as available from Addgene and SEVA repositories.

### Bacterial cultivation

*P. brasiliensis* Ab134 wild type and Δ*pppA* mutant strains were cultivated at 30°C on BD Difco™ Marine Agar 2216 (MA) or in BD Difco™ Marine Broth 2216 (MB) for 18-24 hours unless otherwise noted. Kanamycin (200 µg/mL) was used for mutant selection as appropriate. *E. coli* strains were cultured in BD Difco™ Luria Broth (LB) or on LB agar for 18-20 hours. Kanamycin (50 µg/mL) was used for mutant selection as appropriate. *E. coli* DH5α was used for plasmid vector propagation and *E. coli* S17-1 for conjugation with Ab134. All strains were cryo-preserved in 20% glycerol [*v/v*] at −80 °C.

### Plasmid transfer into *P. brasiliensis* Ab134 by conjugation

#### Plate mating for plasmid transfer

Plasmid vectors were transferred into Ab134 by conjugation from *E. coli* S17-1, using a similar protocol as previously described [3]. Vectors were first electroporated into *E. coli* S17-1.To prepare cells for conjugation, Ab134 was cultured by inoculating 200 µL of overnight seed cultures in 10 mL of MB in a 50 mL falcon tube, and *E. coli* S17-1 was cultured by inoculating 200 µL of overnight seed cultures in 10 mL of LB with kanamycin (50 μg/mL) in a 50 mL falcon tube at 30°C, 200 rpm, until reaching optical density at 600 nm (OD_600_) ∼ 0.4 to 0.6. Cells were then harvested by centrifugation at 5000 rpm for 5 minutes. The cell pellets were resuspended in 1 mL of LB for *E. coli* S17-1 and 1 mL of MB for Ab134. The conjugation mixture of *E. coli* S17-1 and Ab134 was prepared by mixing 500 µL of each concentrated cell suspension, and 500 µL of this mixture was spread evenly onto MA plates followed by a 19-hour incubation at 30 °C. Selection of exconjugants was carried out by streaking a loopful of the mixed cell mass onto MA (20 mL) plates containing kanamycin (200 μg/mL) for the incoming plasmid and carbenicillin (50 μg/mL) to selectively kill *E. coli* S17-1. Positive clones were inoculated into MB (5 mL) containing kanamycin (200 μg/mL) for plasmid extraction after overnight growth at 30°C. Plasmids were confirmed using restriction digestion and gel electrophoresis (**SI Figure S1**).

#### Filter mating for evaluating conjugation efficiency

The filter mating protocol was adapted from a previously described method [39]. Concentrated cell cultures were prepared as described for plate mating. The conjugation mixture of *E. coli* S17-1 and Ab134 was prepared by mixing 50 µL of each concentrated culture, followed by plating on cellular ester membrane filters (Millipore Sigma, 0.45 µm pore size) without spreading and placed on MA. After a 19-hour incubation at 30°C, the biomass from the conjugation mixture or controls was collected thoroughly using a loop and serially diluted in MB. For the mixture of Ab134 and *E. coli* S17-1 containing pSEVA237R_Pem7, dilutions were prepared to 10^-6^ in 1 mL of MB. For the mixture of Ab134 and *E. coli* S17-1 containing pAM4891, dilutions were prepared to 10^-3^ in 1 mL of MB (**SI Figure S3**).

Negative controls were prepared similarly, using either only Ab134 or *E. coli* S17-1 containing each plasmid. Finally, 100 µL of the diluted culture was plated on MA plates containing kanamycin (200 μg/mL) to select for the incoming plasmid and carbenicillin (100 μg/mL) to selectively kill *E. coli* S17-1. Additionally, the Ab134 only culture was serially diluted to 10^-7^ in 1 mL of MB, and 100 µL of the diluted culture was plated on MA to count the total number of Ab134 recipients (**SI Figure S3**). The final dilution was 10^-8^. The assay was performed in triplicates. Colony-forming units (CFU) per mL were counted, and the log_10_ of the conjugation efficiency was calculated using the following equation:

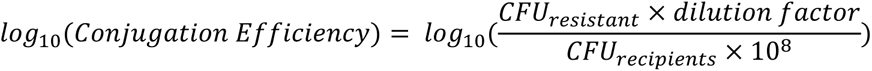

### Plasmid transfer into *P. brasiliensis* Ab134 by electroporation

To prepare Ab134 electro-competent cells, fresh cultures were started by inoculating 200 µL of overnight seed cultures into 10 mL of marine broth in a 50 mL falcon tube. Cultures were incubated at 30°C, 200 rpm, until an OD_600_ between 0.4 and 0.6 was reached. Cells were harvested by centrifugation at 4°C, 5000 rpm for 5 min. Cells were washed three times with 10 mL of ice-cold, filter-sterilized 250 mM sucrose solution [40]. Washed cells were resuspended in the residual sucrose solution, and 100 µL of the concentrated cell suspension were aliquoted for electroporation. Fresh electrocompetent cells were essential for optimal results. For each electroporation, 2 µL (126 ng) of pAM4891 or 2 µL (138.4 ng) of pSEVA237R_Pem7 was added to the cell suspension and mixed gently. The electroporation was carried out in a 0.2 cm ice-cold electroporation cuvette using an electroporator set to 1.8 kV. The expected time constant was around 5.2 – 5.4 ms. Electro-shocked cells were immediately mixed with 500 µL of ice-cold MB and allowed to recover at 30°C with shaking at 200 rpm for 1.5 hours. Recovered cells were harvested by centrifuging at 15,000 rpm for 15 seconds and resuspended in the remaining MB (∼ 100 µL) for plating. MA plates containing kanamycin (200 µg/mL) were used to select positive clones. Competent cells electroporated without plasmid were always used as the control (**SI Figure S4)**. Successful transformants were confirmed by observing fluorescence (**SI Figure S4**). Fluorescence imaging was obtained as described below. Fluorescent colony-forming units (CFU) were counted to calculate electroporation efficiency (CFU per µg plasmid DNA) (**SI Table S4**). The amount of plasmid used was quantified three times by NanoDrop (**SI Table S6**) and confirmed by gel electrophoresis. The assay was performed in triplicates.

### Relative plasmid copy number determination

Total DNA was isolated from *P. brasiliensis* Ab134 carrying one of three plasmids (pAM4891, pSEVA234M, or pSEVA237R_Pem7) using the GenElute^TM^ Bacterial Genomic DNA Kit (Sigma-Aldrich) and following the manufacturer’s instructions. The quality of the total DNA was evaluated by gel electrophoresis and Nanodrop measurements. DNA samples were sequenced using Illumina paired-end whole genome sequencing (400 Mbp, 2.67 M reads package) by the SeqCenter. Sample libraries were prepared using the Illumina DNA Prep kit along with IDT 10bp UDI indices. Sequencing was conducted on an Illumina NextSeq 2000, generating 2×151bp paired-end reads. Demultiplexing, quality control and adapter trimming was performed using bcl-convert (v3.9.3). The obtained sequence reads (**SI Table S1**) were mapped to the reference sequences – Ab134 genome (GenBank: GCA_018282095.1) [3] plus each of the plasmids introduced using the in-built “Map to References” function with medium sensitivity and five iterations in Geneious. In the alignment file, the coverage statistics, which indicates the number of reads mapped at each position, was used to calculate the plasmid copy number as shown below:

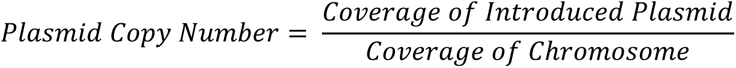

### Growth curves

The growth of each bacterial strain was tracked for 46 h using a microtiter plate method. Seed cultures were prepared by inoculating 50 µL of each cryo-preserved culture into 5 mL of MB. All cultures were incubated at 30°C overnight (18-20 h). Seed cultures were then diluted in MB to OD_600_ of 0.05. OD-standardized cultures were dispensed (200 µL into each well, 10 replicates for each strain) into Falcon™ 96-well, non-treated, flat-bottom microplate using a multi-channel pipettor. Border wells were filled with MB to maintain humidity. Plates were incubated at 30°C and shaken linearly at a speed of 887 revolutions per minute (amplitude 1 mm) using an Infinite M Plex plate reader (Tecan). OD_600_ readings of each well were taken every 30 min over 46 h. A lane containing only MB was used as a blank. The area under the growth curve (not log-scaled) was calculated using the trapezoidal method in the R integrated development environment. Protocols previously described for calculating the lag phase were followed [41]. The OD_600_ values were transformed to ln_(OD600)_, and slopes were calculated for consecutive windows of five time points (e.g., time points 1–5, 2–6, 3–7, etc.) across the entire 46-hour period. The maximum slope among these five-point windows was identified, and five-point slopes that were ≥85% of this maximum slope defined the exponential phase of each growth curve. The first time point with a slope ≥85% of the maximum slope was designated as the end of the lag phase. The R integrated development environment was used to calculate the lag phase for each growth curve.

### Plasmid stability comparison

Plasmid stability in *P. brasiliensis* Ab134 was evaluated for 5 serial passages (120 hours, approximately 50 generations) in the absence of selective pressure. Cultures of Ab134 carrying either pAM4891 or pSEVA237R_Pem7 were initiated by inoculating 50 µL of cryopreserved stocks into 5 mL of marine broth (MB) containing kanamycin (200 µg/mL) and incubating overnight at 30°C, 200 rpm. Cells were then harvested and resuspended in fresh MB to remove kanamycin. An aliquot was serially diluted 1:10^6^ in 1 mL MB and 100 µL was plated on MA plates to assess plasmid retention at the start timepoint T0 with a final dilution of 10^-7^. The first passage was initiated by inoculating 3 µL of the washed overnight seed culture into 3 mL of fresh MB (1:1000 dilution) and incubating at 30°C, 200 rpm, for 24 hours. After each 24-hour incubation, cultures were serially diluted 1:10^7^ in 1 mL of MB, and 100 µL of the diluted culture was plated on MA to count fluorescent colonies, with a final dilution of 10^-8^. This process was repeated for each subsequent 24-hour time point (48, 72, 96, and 120 hours). The final dilution was maintained at 10^-8^, except for Ab134 carrying pSEVA237R_Pem7 at 96 hours (10^-9^) and Ab134 carrying both pAM4891 and pSEVA237R_Pem7 at 120 hours (2 × 10^-8^). The assay was performed in quadruplicates. Plasmid stability was calculated as a percentage using the following equation:

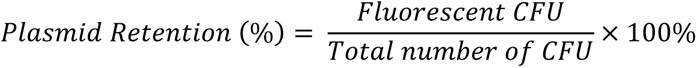

### Swarming motility assays

Swarming assays were performed as previously described [3]. Details are provided here for convenience. Briefly, swarming agar with reduced surface tension was prepared by combining MB and 0.5% (*w/v*) of Eiken agar (Eiken Chemical CO., Japan). The autoclaved agar medium equilibrated to 55 °C for 2 hours was aliquoted onto Petri dishes (exactly 20 mL), which were air dried for 20 min in a laminar flow hood. Fresh cultures of bacterial strains were normalized to OD_600_ of 1.0, and 5 μL of OD-normalized cultures were dropped on the center of each plate. Plates were air dried until the droplet of culture was absorbed and incubated at 30°C. Visible light pictures were taken using a cell phone camera. Fluorescence imaging was obtained as described below. Pictures were taken daily for 3 days.

### Fluorescent imaging

Fluorescent imaging was performed using a ChemiDoc™ MP Imaging System, with Cy2 (532 nm / 28 mm) and Alexa 546 (602 nm / 50 mm) channels for green and red fluorescence, respectively.

## Acknowledgments

We thank Dr. V. de Lorenzo (SEVA collection) for pSEVA234M and pSEVA237R_Pem7, Dr. S. Golden for pAM4891 (Addgene plasmid #120080), and Dr. F. Thompson (Federal University of Rio de Janeiro) and Dr. R. Berlinck (University of São Paulo at São Carlos) for the *P. brasiliensis* Ab134 wild-type strain. Financial support for this work was provided by the Division of Integrative Organismal Systems of the National Science Foundation under grant 1917492 to ASE.

## Author contributions

A.S.E and Y.D. conceptualized the project. Y.D. generated *Pseudovibrio* strains and plasmid related data. A.S.E. advised Y.D. A.S.E. obtained funding for the project. A.S.E. and Y.D. wrote the paper.

## Notes

### Competing Interest Statement

The authors have declared no competing interest.

### Summary of Updates

This version of the manuscript has been revised to follow peer review suggestions. We expanded on genetic tool development for Pseudovibrio and removed sponge studies.

## References

1. Romano S: Ecology and biotechnological potential of bacteria belonging to the genus *Pseudovibrio*. Appl Environ Microbiol 2018, 84(8).

2. Petersen LE, Moeller M, Versluis D, Nietzer S, Kellermann MY, Schupp PJ: Mono- and multispecies biofilms from a crustose coralline alga induce settlement in the scleractinian coral. Coral Reefs 2021, 40(2):381–394.

3. Ioca LP, Dai Y, Kunakom S, Diaz-Espinosa J, Krunic A, Crnkovic CM, Orjala J, Sanchez LM, Ferreira AG, Berlinck RGS et al: A family of nonribosomal peptides modulate collective behavior in *Pseudovibrio* bacteria isolated from marine sponges. Angew Chem Int Ed Engl 2021, 60(29):15891–15898.

4. Verstraeten N, Braeken K, Debkumari B, Fauvart M, Fransaer J, Vermant J, Michiels J: Living on a surface: swarming and biofilm formation. Trends Microbiol 2008, 16(10):496–506.

5. Gutleben J, Loureiro C, Ramirez Romero LA, Shetty S, Wijffels RH, Smidt H, Sipkema D: Cultivation of bacteria from *Aplysina aerophoba*: effects of oxygen and nutrient gradients. Front Microbiol 2020, 11:175.

6. Taton A, Unglaub F, Wright NE, Zeng WY, Paz-Yepes J, Brahamsha B, Palenik B, Peterson TC, Haerizadeh F, Golden SS et al: Broad-host-range vector system for synthetic biology and biotechnology in cyanobacteria. Nucleic Acids Res 2014, 42(17):e136.

7. Silva-Rocha R, Martinez-Garcia E, Calles B, Chavarria M, Arce-Rodriguez A, de Las Heras A, Paez-Espino AD, Durante-Rodriguez G, Kim J, Nikel PI et al: The Standard European Vector Architecture (SEVA): a coherent platform for the analysis and deployment of complex prokaryotic phenotypes. Nucleic Acids Res 2013, 41(Database issue):D666–675.

8. Fernandez HN, Kretsch AM, Kunakom S, Kadjo AE, Mitchell DA, Eustaquio AS: High-yield lasso peptide production in a *Burkholderia* bacterial host by plasmid copy number engineering. ACS Synth Biol 2024, 13(1):337–350.

9. Rouches MV, Xu Y, Cortes LBG, Lambert G: A plasmid system with tunable copy number. Nat Commun 2022, 13(1):3908.

10. Mi J, Sydow A, Schempp F, Becher D, Schewe H, Schrader J, Buchhaupt M: Investigation of plasmid-induced growth defect in *Pseudomonas putida*. J Biotechnol 2016, 231:167–173.

11. Dorado-Morales P, Garcillan-Barcia MP, Lasa I, Solano C: Fitness cost evolution of natural plasmids of *Staphylococcus aureus*. mBio 2021, 12(1).

12. Calos MP: DNA sequence for a low-level promoter of the *lac* repressor gene and an ‘up’ promoter mutation. Nature 1978, 274(5673):762–765.

13. Dong H, Nilsson L, Kurland CG: Gratuitous overexpression of genes in *Escherichia coli* leads to growth inhibition and ribosome destruction. J Bacteriol 1995, 177(6):1497–1504.

14. Brunelle BW, Bearson BL, Bearson SMD, Casey TA: Multidrug-resistant *Salmonella enterica* serovar typhimurium isolates are resistant to antibiotics that influence their swimming and swarming motility. Msphere 2017, 2(6).

15. Dörries K, Schlueter R, Lalk M: Impact of antibiotics with various target sites on the metabolome of *Staphylococcus aureus*. Antimicrob Agents Chemother 2014, 58(12):7151–7163.

16. Shieh WY, Lin YT, Jean WD: *Pseudovibrio denitrificans* gen. nov., sp. nov., a marine, facultatively anaerobic, fermentative bacterium capable of denitrification. Int J Syst Evol Microbiol 2004, 54(Pt 6):2307–2312.

17. Philippe N, Alcaraz JP, Coursange E, Geiselmann J, Schneider D: Improvement of pCVD442, a suicide plasmid for gene allele exchange in bacteria. Plasmid 2004, 51(3):246–255.

18. Meyer R: Replication and conjugative mobilization of broad host-range IncQ plasmids. Plasmid 2009, 62(2):57–70.

19. Priefer UB, Simon R, Puhler A: Extension of the host range of *Escherichia coli* vectors by incorporation of RSF1010 replication and mobilization functions. J Bacteriol 1985, 163(1):324–330.

20. Frey J, Bagdasarian MM, Bagdasarian M: Replication and copy number control of the broad-host-range plasmid RSF1010. Gene 1992, 113(1):101–106.

21. Antoine R, Locht C: Isolation and molecular characterization of a novel broad-host-range plasmid from *Bordetella bronchiseptica* with sequence similarities to plasmids from Gram-positive organisms. Mol Microbiol 1992, 6(13):1785–1799.

22. Chan CH, Levar CE, Zacharoff L, Badalamenti JP, Bond DR: Scarless genome editing and stable inducible expression vectors for *Geobacter sulfurreducens*. Appl Environ Microbiol 2015, 81(20):7178–7186.

23. Lin L, Guo W, Xing Y, Zhang X, Li Z, Hu C, Li S, Li Y, An Q: The actinobacterium *Microbacterium* sp. 16SH accepts pBBR1-based pPROBE vectors, forms biofilms, invades roots, and fixes N_2_ associated with micropropagated sugarcane plants. Appl Microbiol Biotechnol 2012, 93(3):1185–1195.

24. Andrup L, Andersen K: A comparison of the kinetics of plasmid transfer in the conjugation systems encoded by the F plasmid from *Escherichia coli* and plasmid pCF10 from *Enterococcus faecalis*. Microbiology 1999, 145:2001–2009.

25. Ohse M, Takahashi K, Kadowaki Y, Kusaoke H: Effects of plasmid DNA sizes and several other factors on transformation of *Bacillus subtilis* ISW1214 with plasmid DNA by electroporation. Biosci Biotechnol Biochem 1995, 59(8):1433–1437.

26. Szostkova MH, D: The effect of plasmid DNA sizes and other factors on electrotransformation of *Escherichia coli* JM109. Bioelectrochemistry 1998, 47:319–323.

27. Stein DC, Gregoire S, Piekarowicz A: Restriction of plasmid DNA during transformation but not conjugation in *Neisseria gonorrhoeae*. Infect Immun 1988, 56(1):112–116.

28. Read TD, Thomas AT, Wilkins BM: Evasion of type-I and type-II DNA restriction systems by Incl1 plasmid Collb-P9 during transfer by bacterial conjugation. Mol Microbiol 1992, 6(14):1933–1941.

29. Willetts N, Wilkins B: Processing of plasmid DNA during bacterial conjugation. Microbiol Rev 1984, 48(1):24–41.

30. Tesson F, Herve A, Mordret E, Touchon M, d’Humieres C, Cury J, Bernheim A: Systematic and quantitative view of the antiviral arsenal of prokaryotes. Nat Commun 2022, 13(1):2561.

31. Pingoud A, Jeltsch A: Structure and function of type II restriction endonucleases. Nucleic Acids Res 2001, 29(18):3705–3727.

32. Blakesley RW, Dodgson JB, Nes IF, Wells RD: Duplex regions in single-stranded Phi-X174 DNA are cleaved by a restriction endonuclease from *Haemophilus aegyptius*. J Biol Chem 1977, 252(20):7300–7306.

33. Nishigaki K, Kaneko Y, Wakuda H, Husimi Y, Tanaka T: Type-II restriction endonucleases cleave single-stranded DNAs in general. Nucleic Acids Res 1985, 13(16):5747–5760.

34. Shachrai I, Zaslaver A, Alon U, Dekel E: Cost of unneeded proteins in *E. coli* is reduced after several generations in exponential growth. Mol Cell 2010, 38(5):758–767.

35. Scott M, Gunderson CW, Mateescu EM, Zhang Z, Hwa T: Interdependence of cell growth and gene expression: origins and consequences. Science 2010, 330(6007):1099-1102.

36. Klumpp S, Scott M, Pedersen S, Hwa T: Molecular crowding limits translation and cell growth. Proc Natl Acad Sci U S A 2013, 110(42):16754–16759.

37. Kafri M, Metzl-Raz E, Jona G, Barkai N: The cost of protein production. Cell Rep 2016, 14(1):22–31.

38. Dai Y, Lourenzon V, Ioca LP, Al-Smadi D, Arnold L, McIntire I, Berlinck RGS, Eustaquio AS: Pseudovibriamides from *Pseudovibrio* marine sponge bacteria promote flagellar motility via transcriptional modulation. mBio 2024:e0311524.

39. Roberts AP, Mullany P, Wilson M: Gene transfer in bacterial biofilms. Methods Enzymol 2001, 336:60–65.

40. Wang H, Griffiths MW: Mg^2+^-free buffer elevates transformation efficiency of *Vibrio parahaemolyticus* by electroporation. Lett Appl Microbiol 2009, 48(3):349–354.

41. Hall BG, Acar H, Nandipati A, Barlow M: Growth rates made easy. Mol Biol Evol 2014, 31(1):232–238.

