## Supplementary Information for "Evaluation of vectors for gene expression in *Pseudovibrio* marine bacteria"

### Tables

**Table S1. Illumina sequencing results of *P. brasiliensis* Ab134 carrying different plasmids.**

| Plasmid vector name | pAM4891 | pSEVA234M | pSEVA237R_Pem7 |
|---|---|---|---|
| **Replicon** | RSF1010 | pBBR1 | pBBR1 |
| **Total Reads** | 5,029,942 | 3,427,816 | 4,393,640 |
| **Assembled to Reference (%)** | 98.95% | 97.75% | 99.33% |
| **Chromosome Coverage** | 111.9 | 66.2 | 94.4 |
| **Plasmid 1 Coverage** | 122.8 | 74.8 | 103 |
| **Plasmid 2 Coverage** | 117.7 | 82.1 | 98.2 |
| **Plasmid 3 Coverage** | 170 | 124.2 | 160.2 |
| **Plasmid 4 Coverage** | 155.8 | 115.8 | 142.9 |
| **Plasmid 5 Coverage** | 109.3 | 94.9 | 116 |
| **Conjugated Plasmid Coverage** | 5,525.6 | 13,838.3 | 13,957.5 |

**Table S2. Area under the curve and lag phase of *P. brasiliensis* Ab134 carrying different plasmids.**

| Strains | Area under the curve (AUC) | Lag phase (h) |
|---|---|---|
| WT_1 | 37.21 | 2.5 |
| WT_2 | 35.52 | 2.5 |
| WT_3 | 39.51 | 2.5 |
| WT_4 | 38.16 | 2.5 |
| WT_5 | 44.69 | 2.5 |
| WT_6 | 36.24 | 2.5 |
| WT_7 | 34.11 | 2.5 |
| WT_8 | 34.84 | 2.5 |
| WT_9 | 34.10 | 2.5 |
| WT_10 | 34.40 | 2.5 |
| pAM4891_1 | 35.12 | 3.5 |
| pAM4891_2 | 33.76 | 3.5 |
| pAM4891_3 | 33.64 | 3.5 |
| pAM4891_4 | 34.22 | 3.5 |
| pAM4891_5 | 32.45 | 3.5 |
| pAM4891_6 | 33.50 | 3.5 |
| pAM4891_7 | 32.77 | 3.5 |
| pAM4891_8 | 32.97 | 3.5 |
| pAM4891_9 | 32.02 | 3.5 |
| pAM4891_10 | 32.68 | 3.5 |
| pSEVA234M_1 | 16.36 | 2 |
| pSEVA234M_2 | 16.00 | 2.5 |
| pSEVA234M_3 | 15.99 | 2.5 |
| pSEVA234M_4 | 16.40 | 2.5 |
| pSEVA234M_5 | 16.50 | 2.5 |
| pSEVA234M_6 | 15.71 | 2.5 |
| pSEVA234M_7 | 16.44 | 2.5 |
| pSEVA234M_8 | 16.25 | 2.5 |
| pSEVA234M_9 | 16.15 | 2.5 |
| pSEVA234M_10 | 15.87 | 2.5 |
| pSEVA237R_Pem7_1 | 41.08 | 2.5 |
| pSEVA237R_Pem7_2 | 41.00 | 2.5 |
| pSEVA237R_Pem7_3 | 40.98 | 2.5 |
| pSEVA237R_Pem7_4 | 40.50 | 2.5 |
| pSEVA237R_Pem7_5 | 39.62 | 2.5 |
| pSEVA237R_Pem7_6 | 40.34 | 3.0 |
| pSEVA237R_Pem7_7 | 40.56 | 3.0 |
| pSEVA237R_Pem7_8 | 42.18 | 3.0 |
| pSEVA237R_Pem7_9 | 42.13 | 3.0 |
| pSEVA237R_Pem7_10 | 40.33 | 3.0 |

**Table S3. Electroporation efficiency of pAM4891 and pSEVA237R_Pem7 into *P. brasiliensis* Ab134.**

| **Plasmid** | **pAM4891** | **pSEVA237R_Pem7** |
|---|---|---|
| Fluorescent CFU (triplicates) | 383 | 412 |
| | 451 | 306 |
| | 412 | 434 |
| Average CFU | 415 | 384 |
| Amount of plasmid DNA (µg) | 0.126 | 0.138 |
| CFU per µg plasmid DNA | 3,295 ± 271 | 2,776 ± 495 |

**Table S4. Conjugation efficiency of pAM4891 and pSEVA237R_Pem7 into *P. brasiliensis* Ab134 from *E. coli* S17-1.**

<table>
<tr><th>Plasmid</th><th>CFU×mL$^{-1}$ (triplicates)</th><th>Dilution factor</th><th>Conjugation efficiency</th></tr>
<tr><td rowspan="3">pAM4891</td><td>$1.4×10^5$</td><td rowspan="4">$10^4$</td><td rowspan="4">$3.86×10^{-6} ± 3.34×10^{-7}$</td></tr>
<tr><td>$1.2×10^5$</td></tr>
<tr><td>$1.4×10^5$</td></tr>
<tr><td>Average CFU×mL$^{-1}$</td><td>$1.3×10^5 ± 1.2×10^4$</td></tr>
<tr><td rowspan="3">pSEVA237R_Pem7</td><td>$5.0×10^7$</td><td rowspan="4">$10^6$</td><td rowspan="4">$2.22×10^{-3} ± 6.69×10^{-4}$</td></tr>
<tr><td>$9.0×10^7$</td></tr>
<tr><td>$9.0×10^7$</td></tr>
<tr><td>Average CFU×mL$^{-1}$</td><td>$7.7×10^7 ± 2.3×10^7$</td></tr>
<tr><td rowspan="3">Total recipient CFU×mL$^{-1}$<br>*P. brasiliensis* Ab134</td><td>$3.38×10^{10}$</td><td rowspan="4">$10^8$</td><td rowspan="4">---</td></tr>
<tr><td>$3.54×10^{10}$</td></tr>
<tr><td>$3.44×10^{10}$</td></tr>
<tr><td>Average CFU×mL$^{-1}$</td><td>$3.45×10^{10} ± 8.08×10^8$</td></tr>
</table>

**Table S5. Plasmid stability of pAM4891 and pSEVA237R_Pem7 in *P. brasiliensis* Ab134.**

| Time (h) | Plasmid | Type | CFU×mL$^{-1}$ (quadruplicates) | | | | Average ratio | Dilution factor |
|---|---|---|---|---|---|---|---|---|
| T0 | pAM4891 | Glowing | $1.12\times10^{10}$ | $1.00\times10^{10}$ | $8.7\times10^{9}$ | $8.9\times10^{9}$ | 100% | $10^{8}$ |
| | | Total | $1.12\times10^{10}$ | $1.00\times10^{10}$ | $8.7\times10^{9}$ | $8.9\times10^{9}$ | | |
| | | Ratio | 100% | 100% | 100% | 100% | | |
| | pSEVA237R_Pem7 | Glowing | $1.62\times10^{10}$ | $1.28\times10^{10}$ | $1.20\times10^{10}$ | $1.40\times10^{10}$ | 100% | |
| | | Total | $1.62\times10^{10}$ | $1.28\times10^{10}$ | $1.20\times10^{10}$ | $1.40\times10^{10}$ | | |
| | | Ratio | 100% | 100% | 100% | 100% | | |
| T24 | pAM4891 | Glowing | $1.07\times10^{10}$ | $1.55\times10^{10}$ | $1.55\times10^{10}$ | $1.24\times10^{10}$ | 99.38% | $10^{8}$ |
| | | Total | $1.09\times10^{10}$ | $1.55\times10^{10}$ | $1.56\times10^{10}$ | $1.24\times10^{10}$ | | |
| | | Ratio | 98.17% | 100% | 99.36% | 100% | | |
| | pSEVA237R_Pem7 | Glowing | $1.91\times10^{10}$ | $2.19\times10^{10}$ | $1.82\times10^{10}$ | $1.92\times10^{10}$ | 100% | |
| | | Total | $1.91\times10^{10}$ | $2.19\times10^{10}$ | $1.82\times10^{10}$ | $1.92\times10^{10}$ | | |
| | | Ratio | 100% | 100% | 100% | 100% | | |
| T48 | pAM4891 | Glowing | $1.77\times10^{10}$ | $1.26\times10^{10}$ | $1.33\times10^{10}$ | $1.29\times10^{10}$ | 99.62% | $10^{8}$ |
| | | Total | $1.77\times10^{10}$ | $1.26\times10^{10}$ | $1.33\times10^{10}$ | $1.31\times10^{10}$ | | |
| | | Ratio | 100% | 100% | 100% | 98.47% | | |
| | pSEVA237R_Pem7 | Glowing | $2.26\times10^{10}$ | $1.77\times10^{10}$ | $2.42\times10^{10}$ | $2.41\times10^{10}$ | 100% | |
| | | Total | $2.26\times10^{10}$ | $1.77\times10^{10}$ | $2.42\times10^{10}$ | $2.41\times10^{10}$ | | |
| | | Ratio | 100% | 100% | 100% | 100% | | |
| T72 | pAM4891 | Glowing | $1.06\times10^{10}$ | $1.38\times10^{10}$ | $1.41\times10^{10}$ | $1.53\times10^{10}$ | 97.25% | $10^{8}$ |
| | | Total | $1.10\times10^{10}$ | $1.40\times10^{10}$ | $1.45\times10^{10}$ | $1.58\times10^{10}$ | | |
| | | Ratio | 96.36% | 98.57% | 97.24% | 96.84% | | |
| | pSEVA237R_Pem7 | Glowing | $2.94\times10^{10}$ | $2.64\times10^{10}$ | $3.27\times10^{10}$ | $2.93\times10^{10}$ | 100% | |
| | | Total | $2.94\times10^{10}$ | $2.64\times10^{10}$ | $3.27\times10^{10}$ | $2.93\times10^{10}$ | | |
| | | Ratio | 100% | 100% | 100% | 100% | | |
| T96 | pAM4891 | Glowing | $1.78\times10^{10}$ | $1.40\times10^{10}$ | $1.20\times10^{10}$ | $1.33\times10^{10}$ | 85.44% | $10^{8}$ |
| | | Total | $2.03\times10^{10}$ | $1.59\times10^{10}$ | $1.36\times10^{10}$ | $1.71\times10^{10}$ | | |
| | | Ratio | 87.68% | 88.05% | 88.24% | 77.78% | | |
| | pSEVA237R_Pem7 | Glowing | $1.8\times10^{10}$ | $2.2\times10^{10}$ | $1.7\times10^{10}$ | $1.7\times10^{10}$ | 100% | $10^{9}$ |
| | | Total | $1.8\times10^{10}$ | $2.2\times10^{10}$ | $1.7\times10^{10}$ | $1.7\times10^{10}$ | | |
| | | Ratio | 100% | 100% | 100% | 100% | | |
| T120 | pAM4891 | Glowing | $1.40\times10^{10}$ | $1.44\times10^{10}$ | $1.12\times10^{10}$ | $1.12\times10^{10}$ | 76.6% | $2\times10^{8}$ |
| | | Total | $1.72\times10^{10}$ | $1.92\times10^{10}$ | $1.12\times10^{10}$ | $1.12\times10^{10}$ | | |
| | | Ratio | 81.4% | 75% | 65.06% | 84.93% | | |
| | pSEVA237R_Pem7 | Glowing | $2.66\times10^{10}$ | $1.92\times10^{10}$ | $2.50\times10^{10}$ | $1.90\times10^{10}$ | 100% | |
| | | Total | $2.66\times10^{10}$ | $1.92\times10^{10}$ | $2.50\times10^{10}$ | $1.90\times10^{10}$ | | |
| | | Ratio | 100% | 100% | 100% | 100% | | |

**Table S6. Concentration of plasmid pAM4891 and pSEVA237R_Pem7 measured by NanoDrop.**

| Plasmid | Concentration (ng/µL) | Average concentration (ng/µL) | 260 nm/280 nm | 260 nm/230 nm |
|---|---|---|---|---|
| pAM4891 | 126.3 | 126.0 | 1.89 | 2.17 |
| | 126.8 | | 1.89 | 2.16 |
| | 125.0 | | 1.99 | 2.15 |
| pSEVA237R_Pem7 | 70.4 | 69.2 | 1.92 | 2.10 |
| | 69.3 | | 1.98 | 2.13 |
| | 67.8 | | 1.86 | 1.93 |

#### Figures

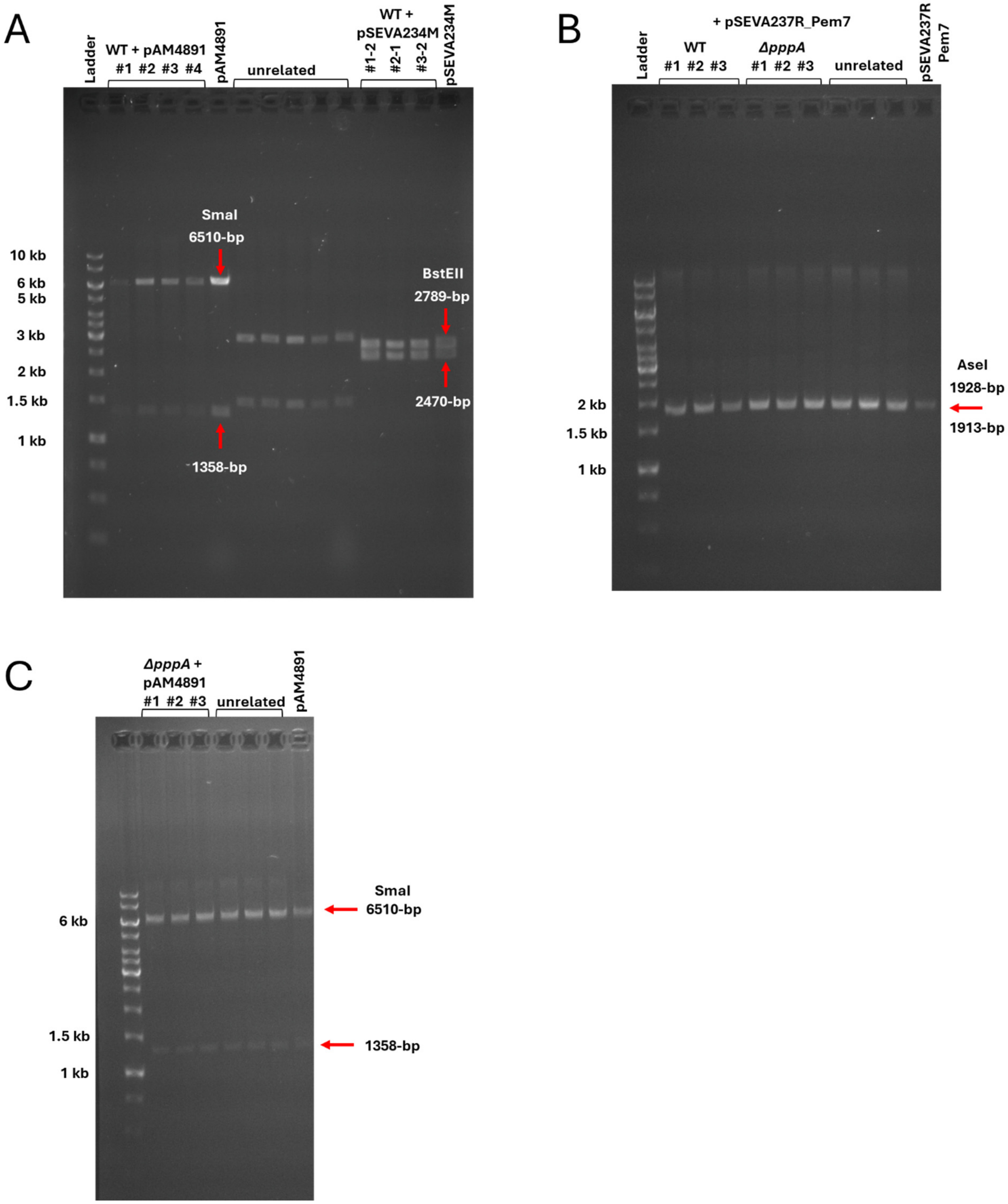

**Figure S1. Plasmid vector DNA isolated from *P. brasiliensis* Ab134.** (**A**) Gel electrophoresis analysis of pAM4891 (RSF1010 replicon) and pSEVA234M (pBBR1) isolated from *P. brasiliensis* Ab134 wild type and digested with SmaI and BstEII respectively. (**B**) Gel electrophoresis analysis of pSEVA237R_Pem7 isolated from *P. brasiliensis* Ab134 wild type and Δ*pppA* mutants. AseI was used for digestion. (**C**) Gel electrophoresis analysis of pAM4891 isolated from *P. brasiliensis* $\Delta$*pppA* mutants. SmaI was used for digestion.

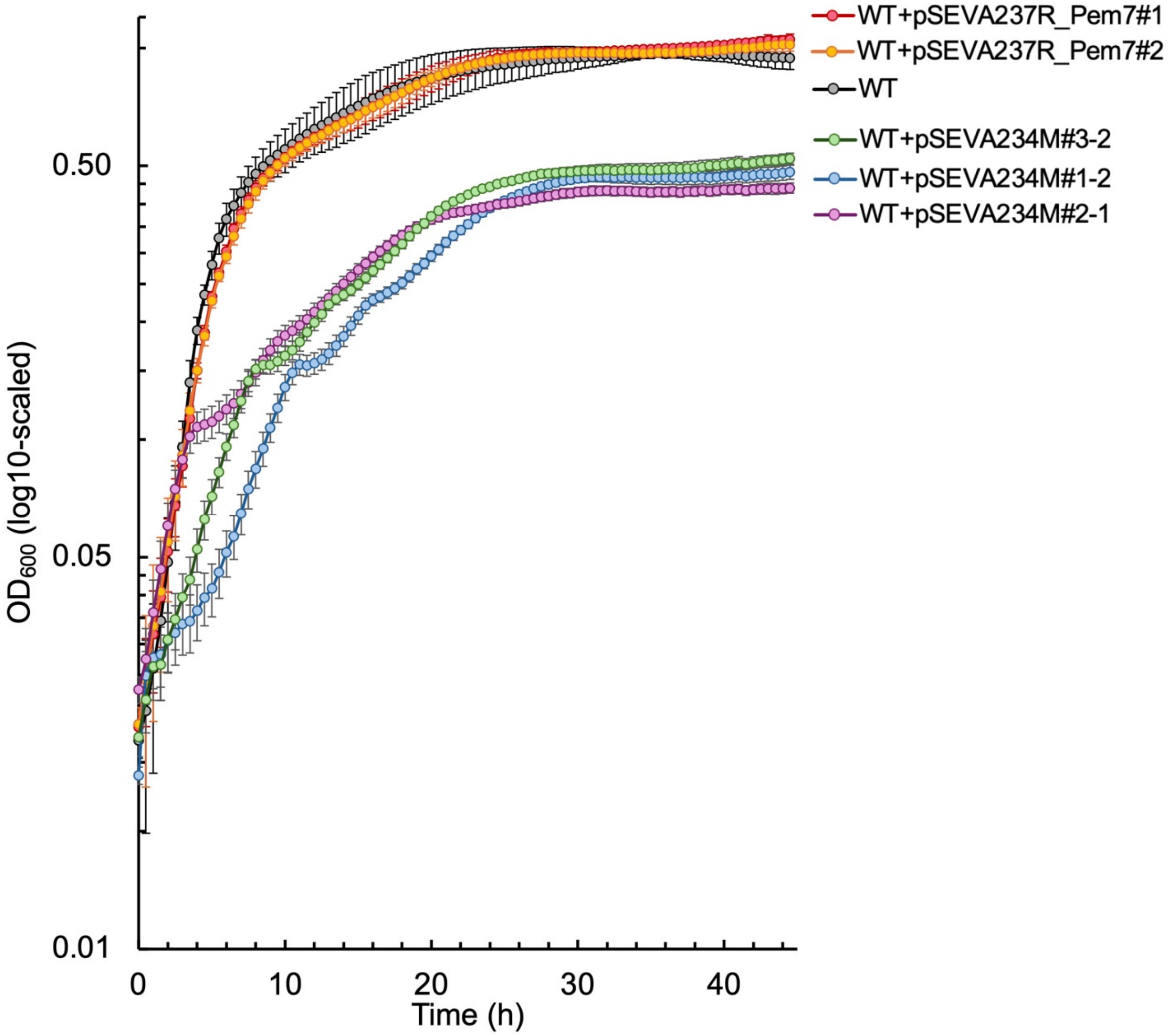

**Figure S2. Growth of independent clones of *P. brasiliensis* Ab134 containing the pBBR1 replicon vectors.** Growth in liquid cultures was measured by $OD_{600}$. *N*=10. Error bars indicate standard deviation.

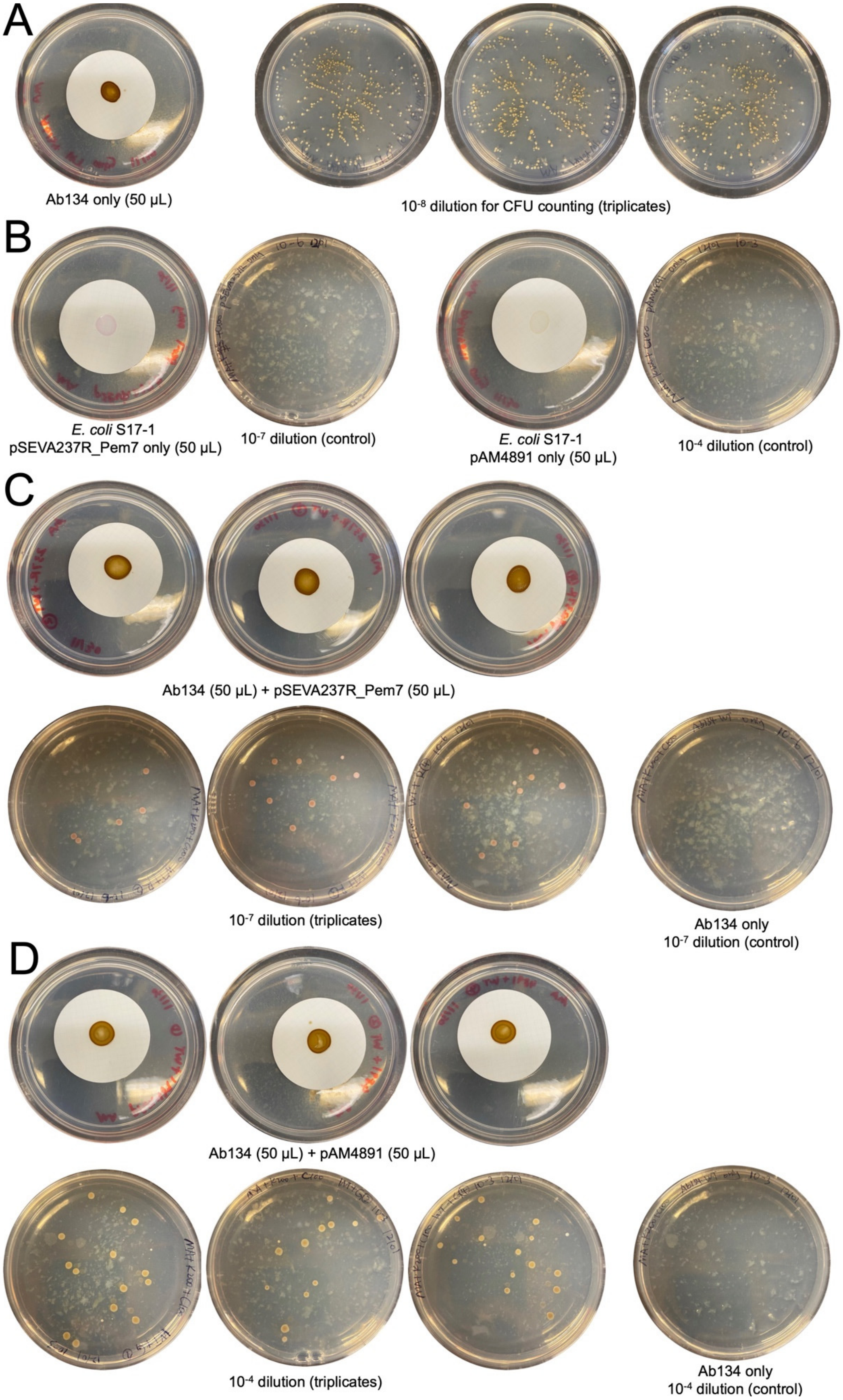
A
Ab134 only (50 µL)
$10^{-8}$ dilution for CFU counting (triplicates)
B
*E. coli* S17-1
pSEVA237R_Pem7 only (50 µL)
$10^{-7}$ dilution (control)
*E. coli* S17-1
pAM4891 only (50 µL)
$10^{-4}$ dilution (control)
C
Ab134 (50 µL) + pSEVA237R_Pem7 (50 µL)
$10^{-7}$ dilution (triplicates)
Ab134 only
$10^{-7}$ dilution (control)
D
Ab134 (50 µL) + pAM4891 (50 µL)
$10^{-4}$ dilution (triplicates)
Ab134 only
$10^{-4}$ dilution (control)

**Figure S3. Filter mating plates for evaluating conjugation efficiency.** (**A**) *Pseudovibrio brasiliensis* Ab134 control on filter paper (left) and triplicate $10^{-8}$ dilutions on marine agar plates without antibiotics for CFU·$mL^{-1}$ counting. (**B**) *Escherichia coli* S17-1 controls carrying either pSEVA237R_Pem7 or pAM4891, with corresponding dilutions on marine agar plates containing kanamycin (200 µg/mL) and carbenicillin (50 µg/mL). (**C**) Conjugation mixtures of *P. brasiliensis* Ab134 and *E. coli* S17-1 carrying pSEVA237R_Pem7 in triplicates, alongside dilutions on marine agar plates containing antibiotics. The $10^{-7}$ dilution plate of *P. brasiliensis* Ab134 only is shown as a control. (**D**) Conjugation mixtures of *P. brasiliensis* Ab134 and *E. coli* S17-1 carrying pAM4891 in triplicates, alongside dilutions on marine agar plates containing antibiotics. The $10^{-4}$ dilution of *P. brasiliensis* Ab134 only is shown as a negative control.

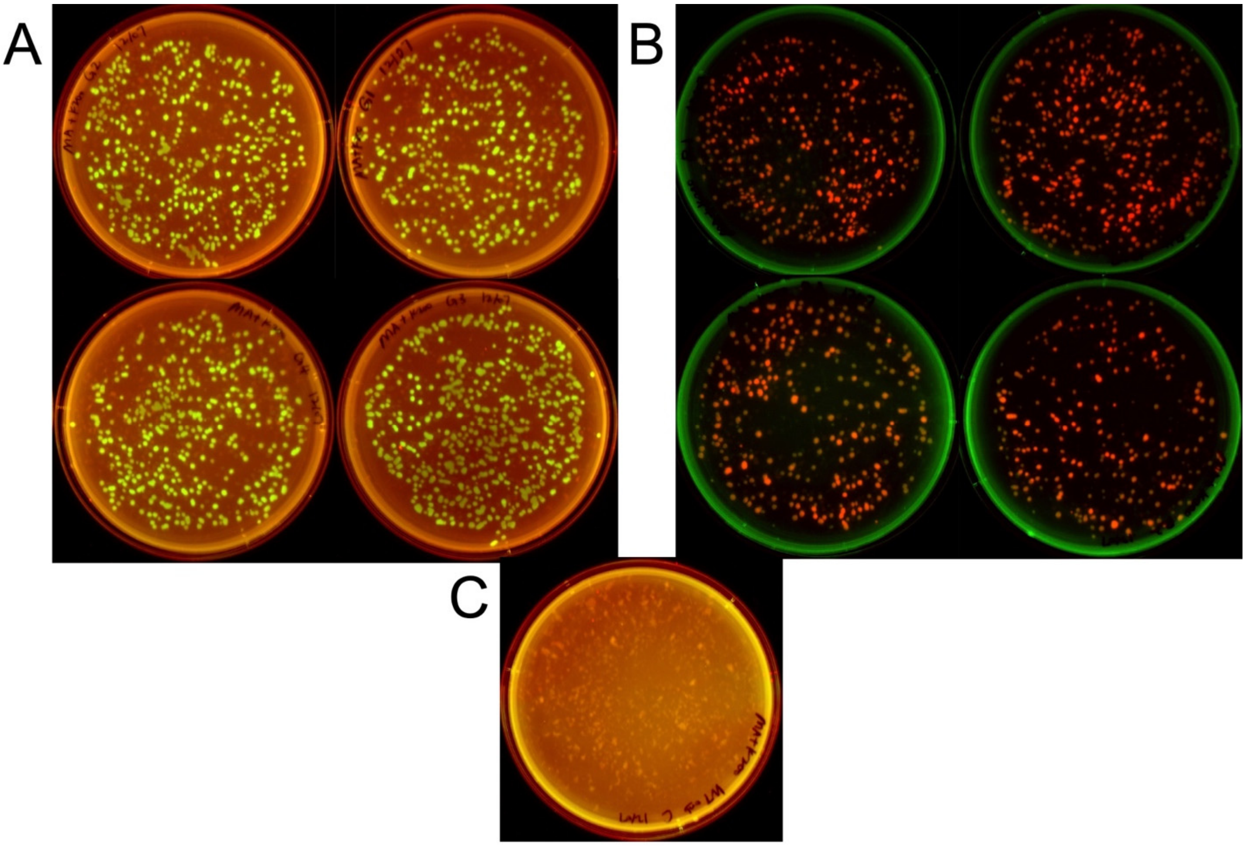

**Figure S4. Fluorescence imaging of quadruplicate images of electroporation results.** (**A**) pAM4891, green fluorescence. (**B**) pSEVA237R_Pem7, red fluorescence. (**C**) Control with no plasmid electroporated. Multi-channel (auto-exposure) with Cy2 (532 nm / 28 mm) and Alexa 546 (602 nm / 50 mm) used to visualize green and red fluorescence, respectively.

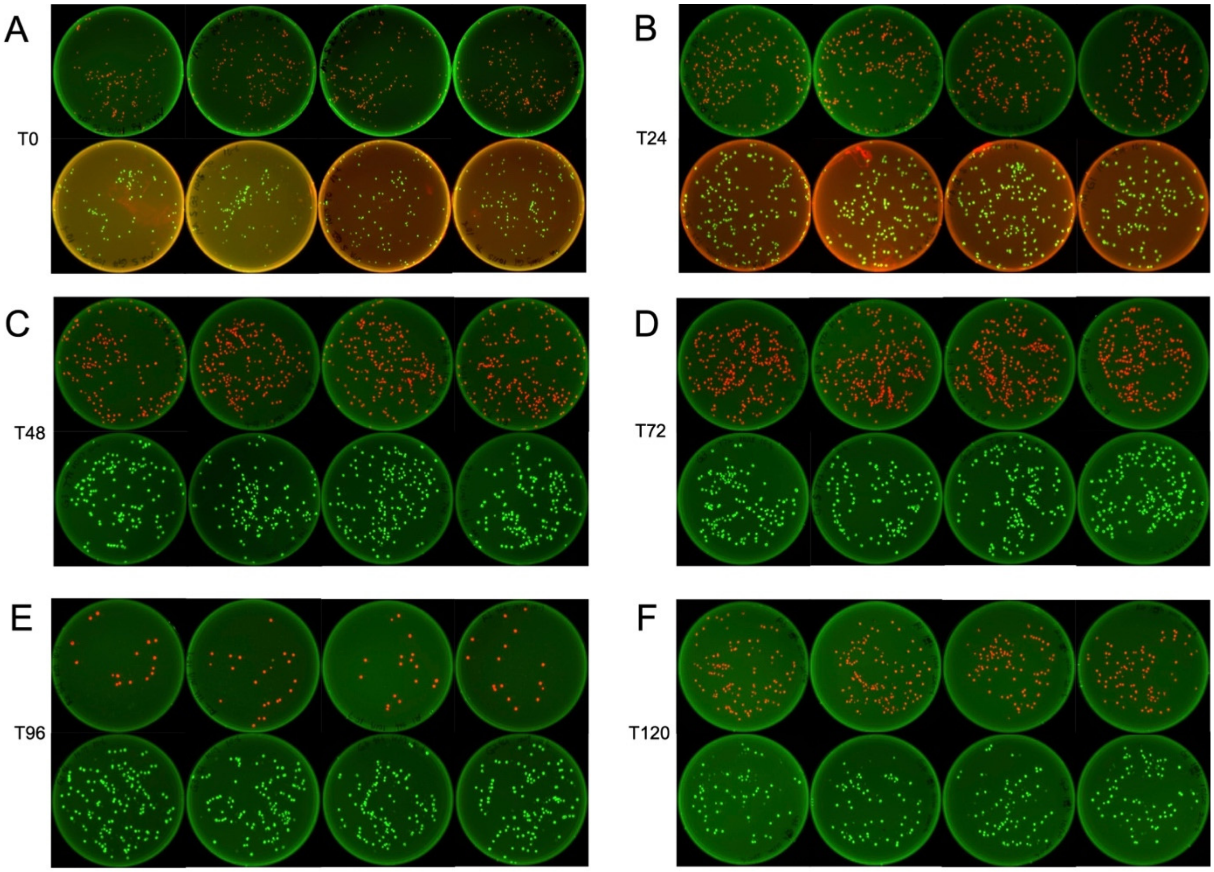

**Figure S5. Fluorescence imaging of quadruplicate plasmid stability results.** Dilution factors are listed in **Table S5**. (**A**) Initial seed culture at T0. (**B**) 1st passage at T24. (**C**) 2nd passage at T48. (**D**) 3rd passage at T72. (**E**) 4th passage at T96. (**F**) 5th passage at T120. pAM4891, green fluorescence; pSEVA237R_Pem7, red fluorescence. Multi-channel (auto-exposure) with Cy2 (532 nm / 28 mm) and Alexa 546 (602 nm / 50 mm) used to visualize green and red fluorescence, respectively.

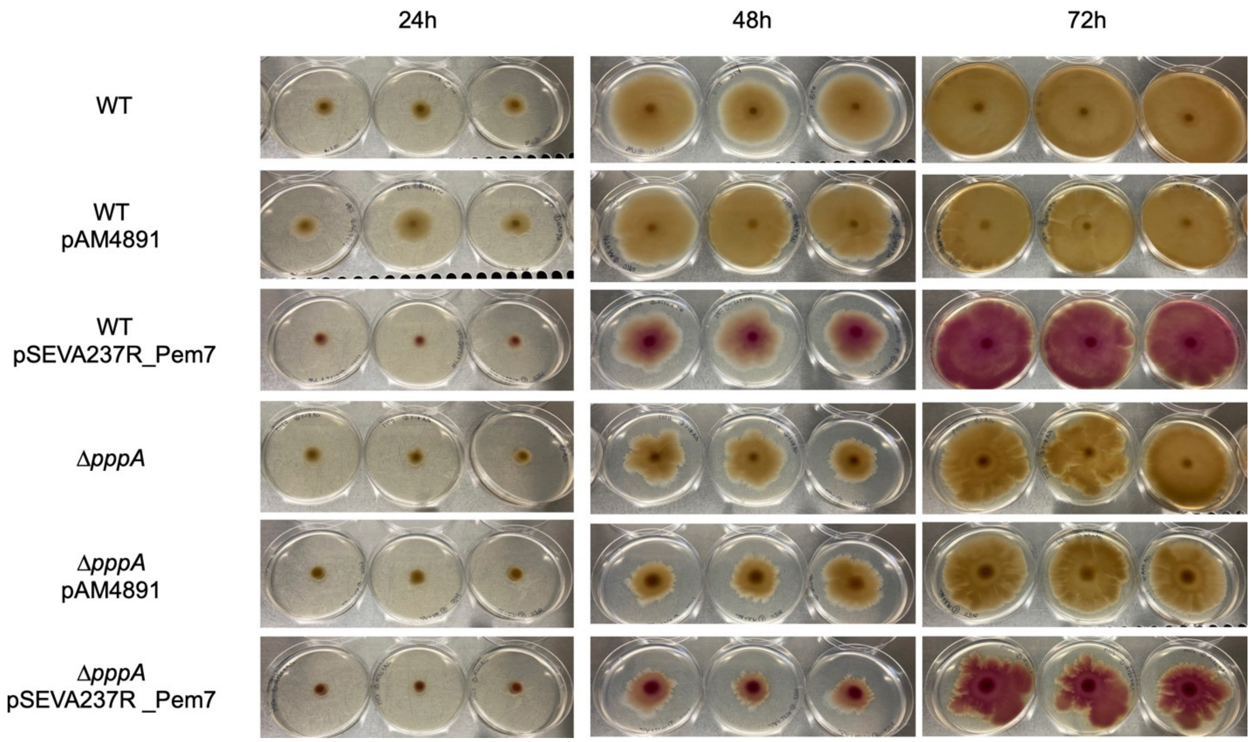

**Figure S6. Triplicate swarming motility assays of *P. brasiliensis* Ab134 wild type or *ΔpppA* mutant compared to strains carrying self-replicative vectors.** Swarming assays performed on marine broth with 0.5 % Eiken agar. Pictures shown were taken at 24, 48, and 72 hours after inoculation, respectively.
